# Diagnostic Labels and Measurement Timing Drive Systematic Inconsistency in ADNI Neuroimaging Data

**DOI:** 10.64898/2026.09.16.752036

**Authors:** Palmer J. Smith, Harry W. Richardson, Katie A. Robertson, Austin J. Dibble, Connor Dalby, Michele Svanera

**Affiliations:** School of Psychology & Neuroscience, University of Glasgow, Glasgow, UK

**Keywords:** ADNI, Label noise, Technical measurement drift, Longitudinal survivor bias, Machine learning for neuroimaging

## Abstract

The Alzheimer’s Disease Neuroimaging Initiative (ADNI) is widely used to train machine learning models for Alzheimer’s disease, yet whether it provides consistent ground truth for predictive modeling has not been systematically tested. In this paper, we showed that ADNI contains three interacting sources of bias with direct implications for machine learning: (a) diagnostic label inconsistency, (b) technical measurement drift, and (c) longitudinal survivor bias. A substantial proportion of cases, particularly within intermediate stages, fall outside ADNI’s own diagnostic thresholds. MRI field strength and evolving processing pipelines introduce significant technical variability in hippocampal volume, while cohort survivor bias arising from differential retention of participants across phases further distorts longitudinal estimates of disease progression. These findings indicated that ADNI does not provide the stable, internally consistent labels often required in machine learning applications. We proposed a practical framework for diagnostic validation, feature harmonization, and cohort accounting, offering guidance for building more robust and biologically meaningful predictive models from large-scale neuroimaging cohorts.

**Graphical Abstract:** Summary of the three interacting sources of bias identified in ADNI (diagnostic label inconsistency, technical measurement drift, and longitudinal survivor bias), their key quantitative findings, and corresponding mitigation strategies for machine learning pipelines.

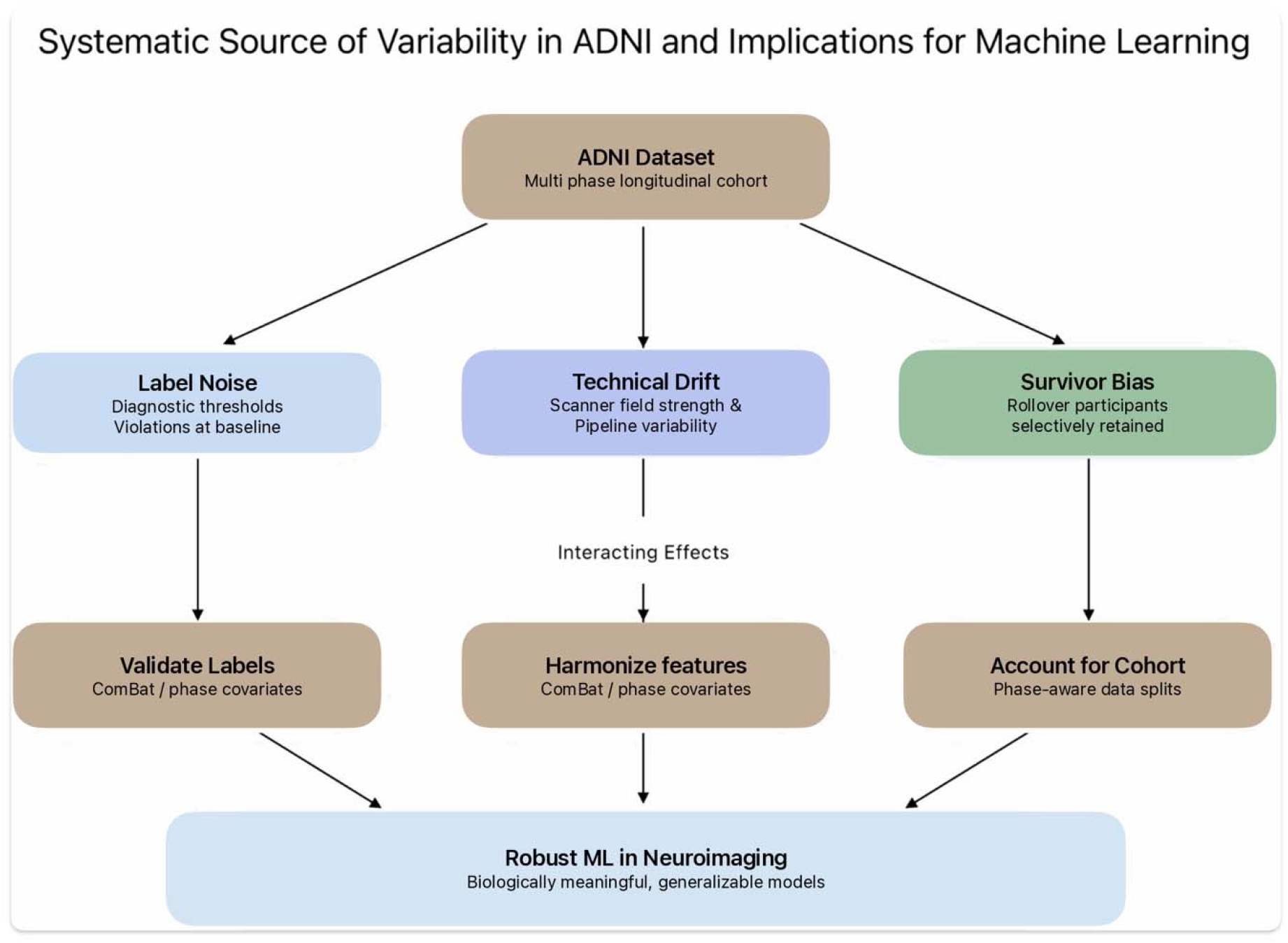

## 1 Introduction

The Alzheimer’s Disease Neuroimaging Initiative (ADNI), established in 2004, is one of the most widely used longitudinal studies in Alzheimer’s disease research (Holbrook et al., 2020; Farina et al., 2023). Across multiple study phases (ADNI1, ADNIGO, ADNI2, ADNI3, ADNI4), ADNI has collected multimodal data including structural MRI, cognitive assessments, and longitudinal clinical follow-up. Owing to its scale, standardization, and public accessibility, ADNI has become a foundational dataset for machine learning applications in neuroimaging, particularly for Alzheimer’s disease diagnostic classification and progression modelling (Bron et al., 2021). This includes recent foundation model approaches aimed at individual brain health estimation (Dibble et al., 2026).

In the context of machine learning, ADNI is frequently treated as a conventional labeled dataset, where its diagnostic categories (defined in Section 3.1) serve as ground truth targets for classification or regression tasks (Petersen et al., 2010). This paradigm underlies a large body of work that treats diagnostic categories as supervised learning targets for predictive modeling using neuroimaging features. However, this widespread usage relies on a critical assumption that ADNI diagnostic labels are internally consistent and closely aligned with the quantitative measurements used in downstream analyses. The data collection study was designed as a longitudinal observational cohort study rather than a machine learning benchmark dataset. Diagnostic assessment occurs during screening visits based on clinical evaluation and cognitive testing, whereas baseline imaging and cognitive measurements may be acquired several weeks later. Another potential issue may rise from cognitive assessments in repeat bias, where subjects who participate across multiple studies (referred to as rollovers in this paper) will get better at taking the test with more attempts (Zheng et al., 2022). If baseline cognitive scores deviate from the thresholds used for diagnostic assignment, the resulting label may not accurately reflect the participant’s measured cognitive state at the time of acquisition. Consequently, ADNI labels could be temporally volatile and cohort-dependent rather than fixed representations of biological state.

Despite these complexities, relatively little attention has been paid to systematically quantifying the extent to which ADNI diagnostic labels align with protocol-defined inclusion criteria, or to evaluating how other structural aspects of the dataset may influence downstream machine learning analyses. This gap is particularly important given that modern machine learning models are highly sensitive to label noise (Frenay & Verleysen, 2014), distribution shifts, and latent confounding factors.

Supervised machine learning methods rest on three implicit assumptions. Diagnostic labels are assumed to accurately reflect the underlying clinical state, feature distributions are assumed to remain sufficiently stable across acquisition settings, and training cohorts are assumed to be broadly representative of the populations to which models will ultimately be applied. In longitudinal neuroimaging datasets such as ADNI, however, these assumptions may be systematically challenged: labels are temporally imprecise for the reasons noted above and may be further compounded by evolving imaging infrastructure and selective participant retention across study phases (Salthouse, 2014). From a computational perspective, these factors correspond to distinct forms of dataset shift, including label noise, acquisition-related covariate shift, and cohort-selection bias (Moreno-Torres et al., 2012). Recent work has begun to address this covariate shift through unified morphological reference spaces that harmonize brain structure measurements across cohorts and diseases (Dalby et al., 2026). Each of which has the potential to affect model training, evaluation, and interpretability.

Motivated by these considerations, we present a systematic investigation of three interacting sources of bias within the ADNI dataset that directly affects machine learning analyses. First, we evaluate diagnostic label consistency by comparing baseline cognitive measurements against protocol-defined thresholds. Second, we quantify technical measurement drift by examining the effects of MRI field strength and software evolution on hippocampal volume estimates across ADNI phases. Third, we assess cohort survivor bias through longitudinal modeling of newly enrolled participants and rollover participants (those continuing from a prior ADNI phase). Together, these analyses characterize how temporal, technical, and cohort-related factors jointly shape the statistical structure of the dataset.

Importantly, rather than treating these analyses as isolated observations, we frame them within a broader methodological perspective: that effective use of ADNI in machine learning requires explicit validation of assumptions at each stage of the modeling pipeline. To this end, we introduce a practical checklist that emerges directly from our findings and is conceptually integrated throughout the paper. This checklist emphasizes (i) verification of diagnostic label consistency, (ii) normalization for technical variability in imaging data, and (iii) accounting for longitudinal cohort effects.

By integrating these considerations into both data preprocessing and model evaluation, we provide actionable guidance for developing more robust and biologically meaningful machine learning models using ADNI. More broadly, we argue that large-scale neuroimaging cohorts should not be treated as static benchmark datasets, but as evolving clinical systems whose structure must be explicitly validated before reliable machine learning inference is possible.

## 2 Methods

This section describes the dataset, analytical framework, and statistical models used to investigate sources of bias within the ADNI cohort. We first detailed the dataset and participant selection criteria, followed by the framework used to evaluate diagnostic label consistency. We then described analyses designed to quantify technical measurement variability and characterize acquisition details across study phases. Finally, we presented a longitudinal modeling approach used to assess cohort survivor effects.

### 2.1 Dataset & Study Design

Data used in this study was obtained from the Alzheimer’s Disease Neuroimaging Initiative (ADNI) database, spanning four completed1 phases (ADNI1, ADNIGO, ADNI2, and ADNI3). It is a multi-site longitudinal study designed to track the progression of Alzheimer’s disease through repeated acquisition of neuroimaging, cognitive, and clinical measures. Recruitment was conducted across more than 50 sites in North America, with each phase enrolling new participants alongside rollover participants from earlier phases. Across these four phases, 3,396 participants were enrolled in total (1,732 female, 1,662 male, 2 unknown; age range 50-89+), contributing an average of 6.82 visits per participant (median 5). Of these, 2,295 unique participants had complete cognitive and structural MRI data and were retained for the present analysis, yielding 8,853 longitudinal observations. Each study visit combines standardized cognitive testing, structural MRI, and at most sites, additional biomarkers such as PET imaging and cerebrospinal fluid measures, though the present analysis focuses on the cognitive and structural MRI data common across all phases.

Participants in ADNI are assigned to diagnostic groups based on clinical evaluation and standardized cognitive testing conducted during a screening visit. These categories include Cognitively Normal (CN), Subjective Memory Complaint (SMC), Early Mild Cognitive Impairment (EMCI), Mild Cognitive Impairment (MCI), Late Mild Cognitive Impairment (LMCI), and Alzheimer’s Disease (AD). These diagnostic groups are defined using combinations of cognitive scores, functional assessments, and clinical judgement, and are intended to represent stages along the Alzheimer’s disease continuum.

Importantly, baseline neuroimaging and cognitive measurements are typically acquired after the screening visit, often with a delay of several weeks. As a result, the measurements used in downstream analyses may not correspond exactly to the values used during diagnostic assignment. In addition, participants may transition between diagnostic categories over time and may continue across multiple ADNI phases through rollover recruitment, introducing further complexity into the dataset structure.

For the present study, we utilized baseline cognitive assessments and structural MRI-derived hippocampal volumes for participants with complete data through CSV files provided by ADNI. Cognitive variables included the Mini-Mental State Examination (MMSE), Clinical Dementia Rating Sum of Boxes (CDR-SB), and Logical Memory II (LM). Hippocampal volume was selected as a representative neuroimaging biomarker due to its established relevance in Alzheimer’s disease progression, with meta-analytic estimates showing annualized atrophy rates roughly three times higher in AD than in cognitively normal older adults (4.6% vs. 1.4%; Barnes et al., 2009). This is consistent with the broader biomarker cascade model of AD progression (Jack et al., 2013).

By focusing on baseline measurements while retaining information about diagnostic assignment and study phase, this dataset enabled a systematic evaluation of consistency between diagnostic labels, quantitative cognitive measures, and imaging-derived biomarkers. This choice was supported by the database’s schedule of assessments, in which the screening MRI is administered concurrently with the MMSE, Clinical Dementia Rating, and Logical Memory I & II (Baseline, M12, M24, etc.). This ensured that the imaging and cognitive data anchoring the baseline diagnostic label were connected at the same visit rather than at a later, potentially discordant timepoint.

### 2.2 Diagnostic Consistency Framework

To evaluate the reliability of diagnostic labels we developed a framework that compares observed cognitive measurements against the protocol-defined thresholds used for participant classification within ADNI.

Each diagnostic category in ADNI is associated with predefined inclusion criteria based on cognitive assessments, including MMSE, CDR-SB, and Logical Memory scores. Using these criteria, permissible ranges (referred to as “safe zones”) for each cognitive variable were acquired directly from ADNI’s established eligibility criteria; the same thresholds used to assign participants to diagnostic categories at enrollment. For example, CN participants are required to score MMSE greater than or equal to 24 and CDR = 0, while MCI participants are required to score MMSE 24-30 and CDR = 0.5, with Logical Memory cutoffs that vary by years of education. Additionally, with ADNI including new diagnostic groups in additional phases (e.g. EMCI, LMCI, etc.) the eligibility range may vary. A participant’s safe zone is therefore the published eligibility range for their assigned diagnostic category, and Out-of-Bounds (OOB) status indicates that a participant’s baseline score falls outside the range.

For each participant, baseline cognitive scores were compared against the corresponding safe zones for their assigned diagnostic category. Participants whose scores fell outside these ranges were classified as OOB, indicating a discrepancy between their measured cognitive status at baseline and the criteria used for diagnostic assignment. It is important to note that OOB classification does not imply protocol violation. Rather, it reflects a temporal mismatch between screening-based diagnostic assignment and baseline measurement. This distinction is critical for interpreting diagnostic labels as potentially noisy or temporally variable targets in machine learning applications.

Logical Memory thresholds were adjusted according to years of education, consistent with ADNI protocol guidelines. This ensured that comparisons account for education-dependent scoring differences.

This framework provides a quantitative basis for assessing the extent to which diagnostic labels can be considered consistent with the cognitive measurements available for model training.

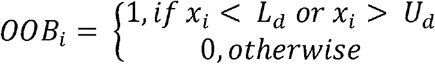

Where *x*_*i*_ represents the participant’s cognitive score and *L*_*d*_, *U*_*d*_ represents the lower and upper bounds for the diagnostic category *d*.

### 2.3 Technical Measurement Drift Analysis

To assess the extent to which imaging infrastructure contributes to variability in neuroimaging biomarkers, we examined the relationship between hippocampal volume and MRI acquisition parameters.

Hippocampal volume was modeled as a function of diagnostic category and MRI field strength (1.5 Tesla vs. 3 Tesla). MRI field strength was treated as a primary variable of interest, as it differs systematically across ADNI phases and has the potential to introduce non-biological variation in measured volumes (Macdonald et al., 2014).

A linear regression model was fit with hippocampal volume *(mm*^*3*^) as the dependent variable, and diagnostic category, MRI field strength, and age as predictors: Hippocampal Volume = *β*_0_ + *β*_1_ (Diagnosis) + *β*_2_ (Field Strength) + *β*_3_ (age) + ε. This specification allowed us to estimate the independent contribution of each predictor while controlling for the others. The AD group scanned at 1.5T was used as the reference category, allowing all other coefficients to be interpreted relative to this baseline.

### 2.4 Scanner Infrastructure Across ADNI Phases

To contextualize the results of the technical measurement analysis, we characterized the distribution of MRI hardware and image processing pipelines across ADNI phases.

For each phase, we summarized the proportion of scans acquired at different field strengths (1.5T and 3T), as well as the primary versions of the FreeSurfer segmentation software (Versions 4.3, 5.1, 6.0; Fischl 2012), whose volumetric outputs are not directly comparable across major releases (Bigler et al., 2020). These summaries provide insight into systematic differences in data acquisition and processing that may influence downstream analyses. More recently, segmentation networks have been proposed that are explicitly trained to be robust to scanner and site variability, offering an alternative to conventional pipeline-based approaches (Svanera et al., 2024).

Because both hardware and software evolved across ADNI phases, this characterization allows us to interpret volumetric differences considering underlying technological changes. It provides a basis for understanding how shifts in scanner field strength and segmentation algorithms may contribute to apparent differences in brain structure measurements across cohorts.

### 2.5 Longitudinal Atrophy Model

To investigate potential cohort survivor bias, we analyzed longitudinal changes in hippocampal volume using a linear mixed-effects modeling framework.

Hippocampal volume was modeled as a function of time since baseline, participant status (newly enrolled vs. rollover participant), and the interaction between these variables. Rollover participants were defined as individuals who continued into a subsequent ADNI phase after initial enrollment, while newly enrolled participants were those entering the study for the first time within a given phase.

Random intercepts and slopes were included for each participant to account for repeated measurements and inter-individual variability. Model parameters were estimated using restricted maximum likelihood (REML) rather than full maximum likelihood, as REML produces unbiased estimates of variance components in mixed-effects models with unbalanced longitudinal data — a property particularly relevant here given that participants contribute unequal numbers of follow-up visits across ADNI phases

This modeling approach enables comparison of both baseline hippocampal volume and longitudinal atrophy rates between groups, a quantitative framework for assessing whether participants who remain in the study differ systematically renfrom newly recruited individuals.

By explicitly modeling these differences, this analysis addresses an important source of bias in longitudinal neuroimaging datasets and highlights the need to account for cohort structure when developing predictive models.

For all models reported in this study, statistical significance was assessed at *a =* 0.05, and exact p-values are reported throughout except where p < 0.001.

## 3 Results

### 3.1 Diagnostic Threshold Violations

Across all ADNI phases, a substantial proportion of participants exhibited baseline cognitive scores that fell outside the protocol-defined thresholds associated with their assigned diagnostic categories (Table 1 below). Violation rates were relatively low in ADNI1, where no rollover cohort existed to introduce temporal drift but increased sharply in later phases. This is particularly true within intermediate diagnostic groups reassessed at each phase’s own entry visit rather than at original enrollment.

**Table 1.**
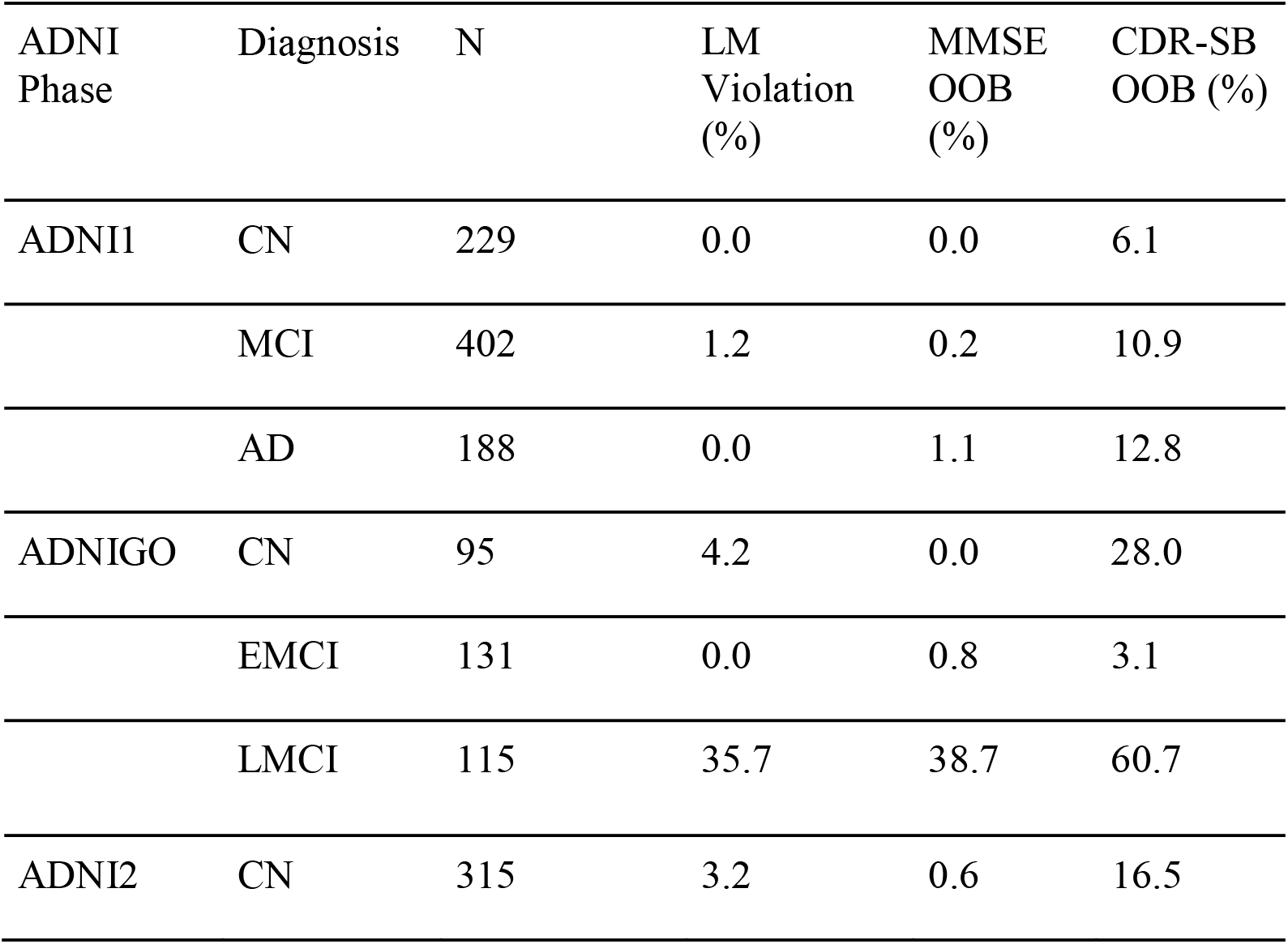
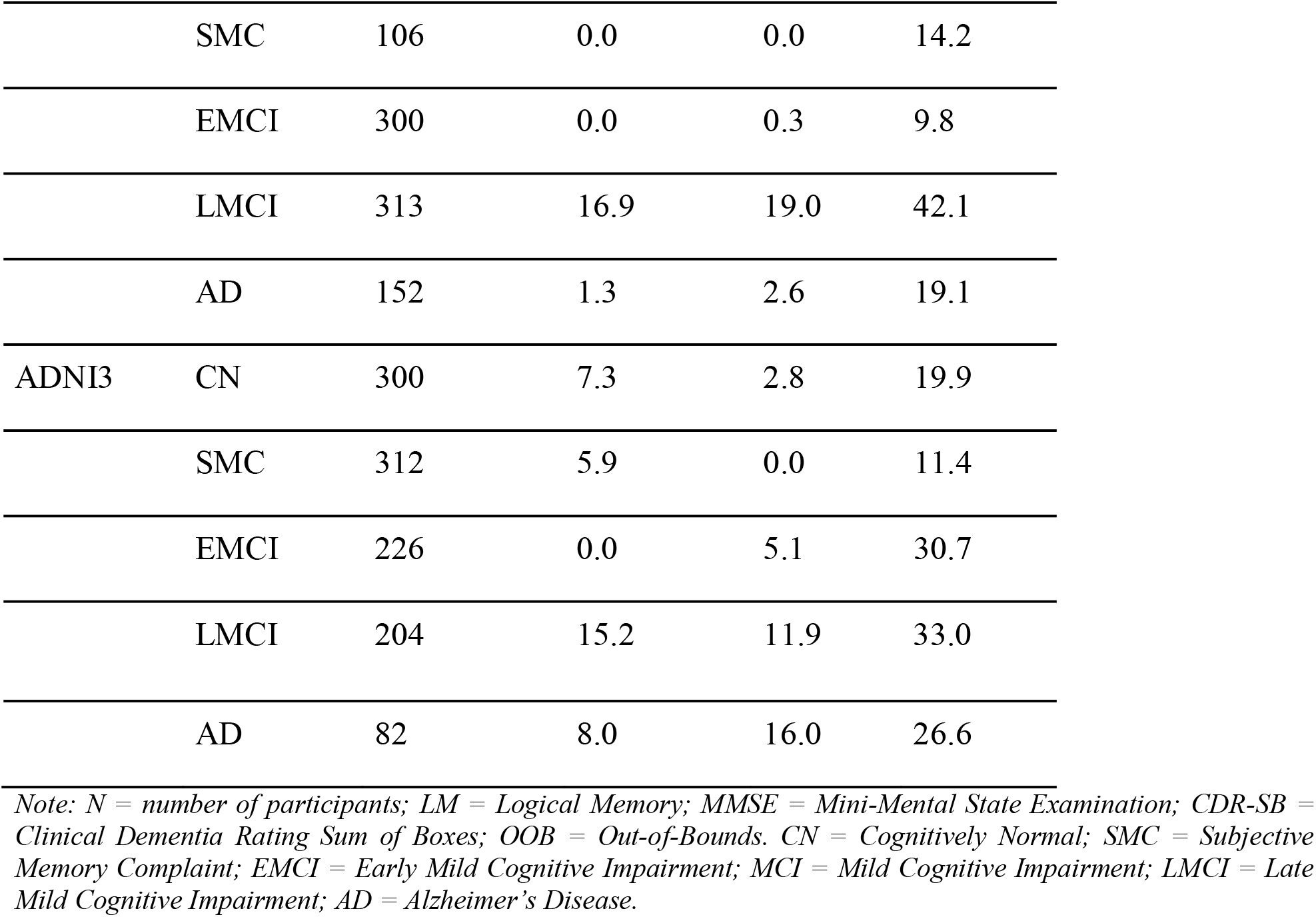

The highest rates of deviation were observed in Early and Late Mild Cognitive Impairments (EMCI & LMCI). Combined across all three cognitive measures, 30.2% of EMCI/LMCI participants fell outside of at least one safe zone, compared to 16.8% of CN/SMC participants and 22% of AD participants. The effect was most pronounced among rollover participants: in the ADNIGO cohort, 89.6% of LMCI participants, those continuing from ADNI and reassessed 3-5 years after their original baseline, fell outside of at least one cognitive safe zone.

Similar patterns were observed in ADNI2 & ADNI3, where intermediate groups consistently showed higher rates of inconsistency than CN & AD groups. To test competing explanations for this pattern, we compared the explanatory power of screening-to-baseline interval length against diagnostic category in predicting out-of-bounds status. Diagnostic category explained substantially more variance in out-of-bounds status (pseudo *R*^2^=*0* 0.035) than interval length, which was not a statistically significant predictor (p = 0.08) once diagnostic category was accounted for. This indicates that these deviations are driven primarily by the structure of diagnostic assignment rather than by short delays between screening and baseline measurement.

### 3.2 Diagnostic Migration Across Phases

Longitudinal transitions between diagnostic categories were visualized using a Sankey diagram illustrating participant movement across ADNI phases (Figure 1). The largest flows were observed within the Mild Cognitive Impairment spectrum, particularly between EMCI and LMCI groups, indicating that these categories represent dynamically evolving clinical states.

**Figure 1.**
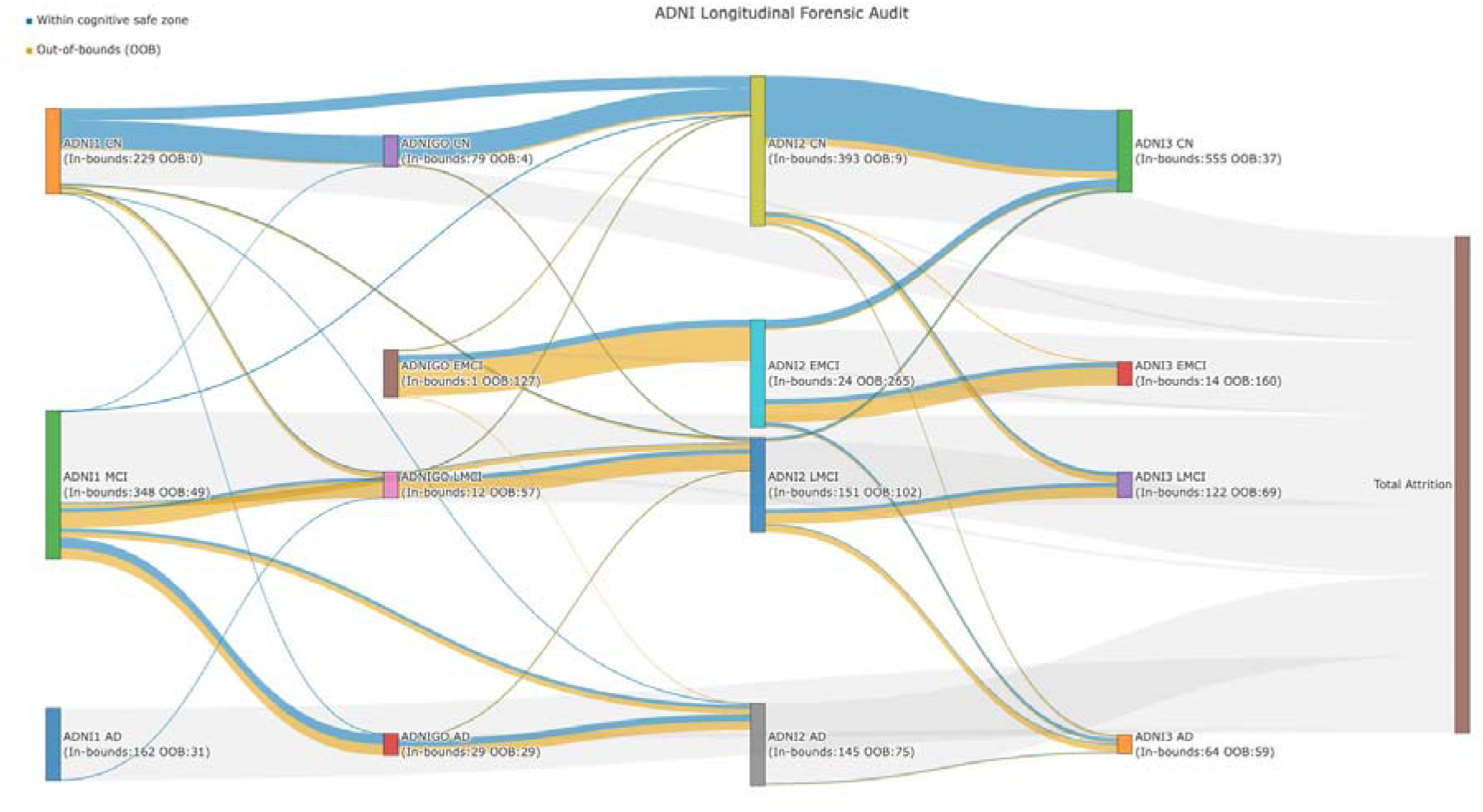
Longitudinal diagnostic transitions across ADNI phases. Each column represents one ADNI phase (ADNI1, ADNIGO, ADNI2, ADNI3, left to right); each block within a column represents one diagnostic category, where block height is proportional to the number of participants in that category at that phase. Connecting paths show individual participants’ transitions between diagnostic categories across consecutive phases, with path width proportional to the number of participants following that ribbon. Path color indicates whether the participant’s cognitive scores fell within (blue) or outside (orange) the protocol-defined safe zone for their assigned category at that phase, as defined in Section 3.2. Paths terminating at “Total Attrition” represent participants who did not continue into a subsequent phase. Node labels are displayed horizontally throughout, as Plotly’s Sankey implementation does not natively support rotated node text.

In contrast, CN and AD groups showed greater stability, with fewer transitions across diagnostic categories. Some transitions that appeared counterintuitive, such as apparent movement from AD to less severe categories, were substantially rarer than they may have first appeared: of 992 total phase-to-phase transitions, only seventy-six (7.7%) represented a decrease in diagnostic severity, and AD→CN transitions account for just two (0.2%). The more common severity-decreasing transitions were LMCI→CN (n=27) and EMCI→CN (n=26), a pattern broadly consistent with documented rates of MCI-to-normal reversion in the clinical literature (Canevelli et al., 2016) rather than a dataset artifact. Apparent long-range crossings in the diagram itself were partly a visual consequence of the fixed vertical node ordering used to keep diagnostic category aligned across phase columns, rather than evidence of additional reversal flows.

Taken together, these results suggest that diagnostic labels in ADNI should be interpreted as temporally dependent cohort descriptors rather than fixed disease states. For machine learning applications, this reinforced the label uncertainty and temporal context when defining prediction targets.

### 3.3 Technical Measurement Drift

Linear regression analysis revealed a strong association between MRI field strength and measured hippocampal volume (Table 2). After controlling for diagnostic group and age, scans acquired at 3T were associated with an average increase of approximately 343 *mm*^3^ in hippocampal volume relative to 1.5T scans.

**Table 2.**

| Predictor Variable | Coefficient ($\beta$) | Std.Error | t-value | p-value |
| --- | --- | --- | --- | --- |
| Intercept | 6,751.00 | 315.40 | 21.40 | < 0.001 |
| Diagnosis: CN | 1,678.00 | 62.68 | 26.77 | < 0.001 |
| Diagnosis: EMCI | 1,226.00 | 79.64 | 15.39 | < 0.001 |
| Diagnosis: LMCI | 833.30 | 82.74 | 10.07 | < 0.001 |
| Diagnosis: MCI | 804.90 | 73.37 | 10.97 | < 0.001 |
| Field Strength (3T vs. 1.5T) | 343.10 | 55.81 | 6.15 | < 0.001 |
| Age (Years) | -58.02 | 3.14 | -18.49 | < 0.001 |

*Note: Table 2's coefficient is mm<sup>3</sup>.*

Diagnostic group differences followed the expected biological gradient, with higher hippocampal volumes observed in CN, EMCI, and LMCI participants, respectively, compared to AD. However, the magnitude of the field strength effect was substantial relative to these group differences, indicating that acquisition parameters contributed significantly to observed volumetric variation.

This result demonstrated that hippocampal volume measurements in ADNI were not solely determined by underlying neurobiology but were also influenced by technical factors related to imaging infrastructure. For machine learning models trained on cross-sectional imaging data, this introduces a risk of learning scanner-specific artifacts rather than clinically meaningful patterns.

These findings highlight the necessity of incorporating acquisition-related variables into model design, either through covariate adjustment or harmonization techniques such as ComBat (Johnson et al., 2007;Fortin et al., 2018; Pomponio et al., 2020).

### 3.4 Scanner Infrastructure Across ADNI Phases

The distribution of MRI hardware and segmentation software varied substantially across ADNI phases (Table 3). ADNI1 relied exclusively on 1.5T scanners, whereas later phases progressively transitioned toward 3T imaging, with ADNI3 using 3T scanners exclusively.

**Table 3.**
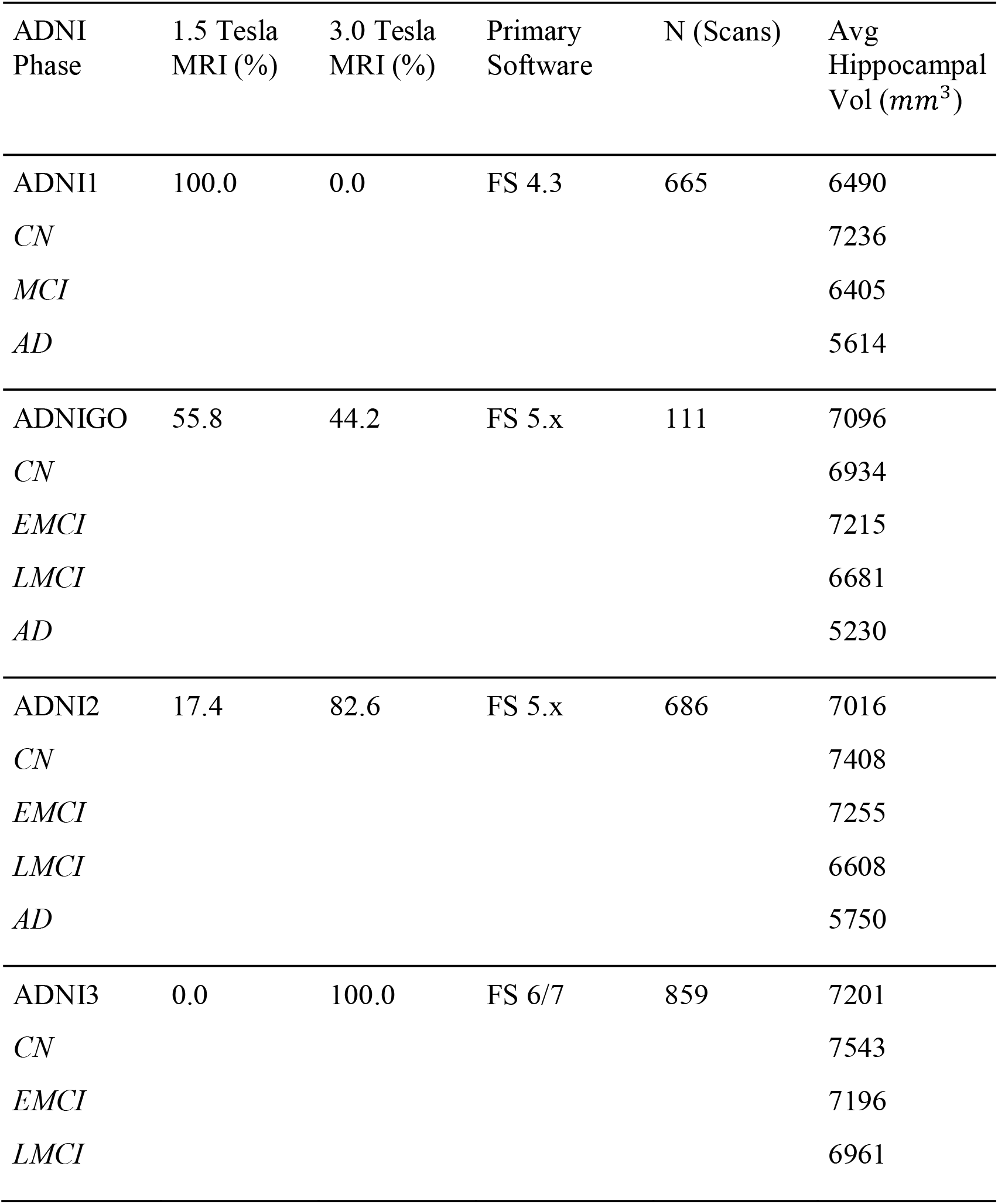
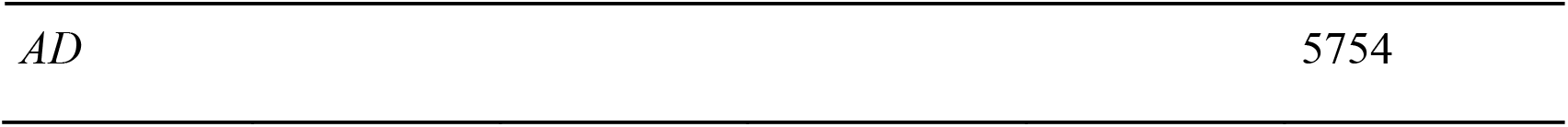

At the same time, segmentation pipelines evolved from earlier versions of FreeSurfer (e.g. version 4.3) to more recent releases (versions 6 & 7). Despite the expectation that newer software versions may produce more conservative volumetric estimates (Schmidt et al., 2018), average hippocampal volumes increased across successive ADNI phases.

This apparent contradiction can be explained by the strong effect of scanner field strength, which appeared to offset or exceed software-related differences. As a result, comparisons of hippocampal volume across ADNI phases reflect a combination of biological variation and systematic differences in acquisition and processing.

For machine learning applications, this implies that pooling data across phases without accounting for these differences may introduce distribution shifts that confound model training and evaluation.

### 3.5 Cohort Survivor Bias

A linear mixed-effects model revealed systematic differences between newly enrolled participants and those who continued across multiple ADNI phases (Table 4). Newly enrolled participants exhibited an estimated baseline hippocampal volume of approximately 6774 *mm*^*3*^and an average decline of 144.95 *mm*^*3*^ per year.

**Table 4.**

| Predictor Variable | Coefficient ($\beta$) | Std.Error | t-value | p-value |
| --- | --- | --- | --- | --- |
| Intercept | 6774.04 | 29.61 | 228.79 | < 0.001 |
| Yrs from Baseline | -144.95 | 4.42 | -32.77 | < 0.001 |
| Status (Rollover) | 425.56 | 52.73 | 8.07 | < 0.001 |
| Interaction (Yrs x Rollover) | 27.87 | 5.68 | 4.90 | < 0.001 |

*Note: Random effects included per-subject intercepts and slopes. $N = 8,853$ observations across 2,295 participants.*

In contrast, rollover participants entered subsequent phases with significantly higher baseline hippocampal volumes, with an estimated advantage of approximately 425 *mm*^*3*^. In addition, these participants exhibited a slower rate of atrophy over time, as indicated by a significant interaction between time and rollover status.

To formally test whether this pattern reflected cohort survivorship rather than compositional differences between groups, we fit an additional linear mixed-effects model predicting hippocampal volume from time since baseline, rollover status, and their interaction, adjusting for age, sex, and diagnosis at enrollment. The interaction between time and rollover status remained significant after adjustment (*β*= (26.8*mm*^3)/year, t = 4.74; N = 8,840 observations, 2,284 participants), confirming that the divergence in decline rates is not attributable to demographic or diagnostic differences between groups (Figure 2). The estimated baseline advantage for rollover participants was substantially attenuated after adjustment *(β* = 117.2, t = 2.76), consistent with rollover status being partly a function of enrolling a less cognitively impaired subset of the cohort to begin with.

**Figure 2.**
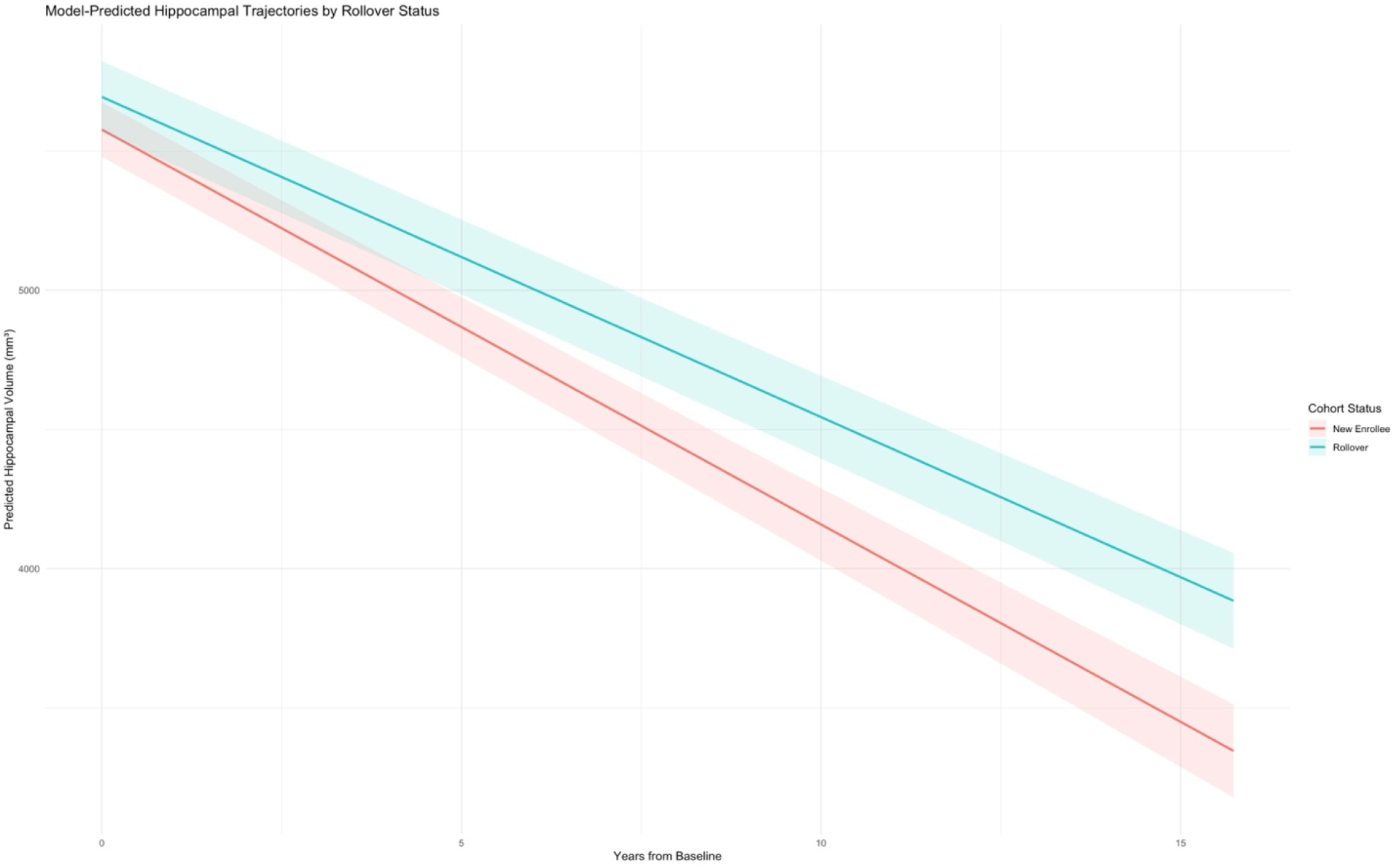
This model displays the progressive decay of hippocampal volume of new participants & participants from previous phases (or rollover participants). See the red line as the new participant populations & the blue as the rollover.

For machine learning models trained on longitudinal data, this effect may lead to systematic underestimation of disease progression rates. Models that do not account for cohort structure may therefore fail to generalize to real-world clinical populations, where more aggressive disease trajectories are present.

Together, these results demonstrate that ADNI contains multiple interacting sources of bias that must be explicitly addressed when developing predictive models.

## 4 Discussion

### 4.1 Conceptual Overview

In this study, we conducted a systematic evaluation of structural properties of the ADNI dataset that have direct implications for machine learning applications. Rather than treating diagnostic labels and imaging measurements as fixed and internally consistent entities, our analyses reveal that ADNI should be understood as a dynamic system shaped by temporal, technical, and cohort-related factors.

Specifically, we identified three interacting sources of bias; (i) variability between diagnostic labels and protocol-defined cognitive thresholds, (ii) systematic measurement differences arising from imaging infrastructure, and (iii) cohort selection effects driven by longitudinal participant retention. Importantly, these factors do not operate in isolation. Instead, they jointly influence the statistical structure of the dataset, shaping both the inputs and targets used in machine learning models.

This perspective challenges the common practice of treating ADNI as a conventional labeled dataset and instead motivates a more cautious and structured approach to its use in predictive modeling.

### 4.2 Diagnostic Labels as Noisy Targets

Our analysis shows that a substantial proportion of participants exhibit cognitive scores that fall outside the thresholds associated with their assigned diagnostic categories. This finding suggests that diagnostic labels in ADNI do not consistently represent the measured cognitive state at baseline.

To test whether OOB status primarily reflects temporal misalignment between screening and baseline or reflects the diagnostic structure itself, we compared the explanatory power of screening-to-baseline gap length against diagnostic category. Gap length was a significant predictor (p= 0.006) but explained a small fraction of variance in OOB status (pseudo *R*^*2*^ *=* 0.003). Diagnostic category alone explained roughly nine times more (pseudo *R*^*2*^*=* 0.028), and this effect was not attenuated by controlling for gap length. This indicates that OOB status is driven predominantly by diagnostic category rather than by the interval between screening and baseline measurement, consistent with ADNI’s diagnostic assignment process that incorporates clinical judgement beyond fixed numeric thresholds.

A key contributing factor is the temporal separation between screening-based diagnostic assignment and baseline data acquisition. Because cognitive performance can fluctuate over relatively short timescales, baseline measurements may diverge from the values used for classification. In addition, diagnostic categories in ADNI are not purely mechanistic thresholds, but incorporate clinical judgment and are designed to support cohort organization within a longitudinal study.

For supervised learning, this means diagnostic category should be treated as a noisy proxy rather than ground truth. In practice, models should be evaluated for sensitivity to label definitions, and reported performance metrics should be stratified by the OOB rate of the training set.

Furthermore, intermediate diagnostic groups such as EMCI and LMCI warrant particular scrutiny given their higher rates of threshold inconsistency.

### 4.3 Imaging Measurements as Non-Stationary Signals

In addition to label variability, hippocampal volume measurements are significantly influenced by imaging infrastructure. This is because ADNI spans multiple study phases with evolving hardware and software, and imaging features reflect a combination of biological variation and acquisition-related effects that cannot be assumed stationary across the dataset.

Consequently, comparisons of imaging-derived biomarkers across ADNI phases must be interpreted in the context of evolving acquisition hardware and processing pipelines. Harmonization methods including ComBat (Fortin et al., 2018; Pomponio et al., 2020) and its longitudinal extension (Beer et al., 2020) provide practical frameworks for correcting such acquisition-related variability and are recommended as standard preprocessing steps when pooling data across ADNI phases.

### 4.4 Integrated Implications for Machine Learning

Diagnostic variability introduces noise in supervision, technical factors induce shifts in feature distributions, and cohort dynamics alter the representation of disease trajectories. Each of these factors has direct consequences for how machine learning models are trained, evaluated and generalized.

These effects highlight the need for a structured approach to model development that explicitly accounts for dataset characteristics. Rather than treating preprocessing and data validation as secondary considerations, our results suggest that they should be central components of the modeling pipeline.

In this context, the analyses presented in this study can be understood as defining a set of practical principles for working with ADNI data. These include verifying the consistency of diagnostic labels with underlying measurements, adjusting for technical variability in imaging features through harmonization (Fortin et al., 2018; Pomponio et al., 2020; Beer et al., 2020), and accounting for cohort composition in longitudinal analyses.

Left unaddressed, these biases have concrete consequences for model performance: label inconsistency produces noisy supervision concentrated in the clinically most ambiguous groups (EMCI/LMCI); technical drift causes models to learn scanner-specific signatures rather than biology, reducing cross-site generalizability; and survivor bias leads to systematic underestimation of disease progression rates, with poor transfer to real-world clinical populations. The graphical abstract summarizes each source of bias alongside its corresponding mitigation strategy.

By integrating these considerations into model design and evaluation, researchers can reduce the risk of learning dataset-specific artifacts and improve the likelihood that their models capture biologically meaningful patterns. More broadly, this framework underscores the importance of critically examining the structure of large-scale neuroimaging datasets when applying machine learning methods, particularly in settings where diagnostic certainty cannot be assumed and data collection processes evolve over time.

Taken together, these findings illustrate a general challenge for computational neuroimaging: supervised learning pipelines implicitly assume label fidelity, feature stationarity, and representative sampling — assumptions that longitudinal cohorts such as ADNI do not automatically satisfy. We provide a starting point for validating these assumptions before model training, and recommend that it is treated as a standard preprocessing step alongside, rather than a substitute for, dataset-specific quality control.

### 4.5 Limitations

Several limitations should be considered when interpreting the findings of this study, particularly in relation to the scope of the analyses and the structure of the ADNI dataset.

First, the present analysis provides evidence of cohort survivor bias but does not explicitly model the underlying mechanisms driving participant retention, such as clinical progression, comorbidities, or site-specific differences. A more detailed characterization of these factors would clarify the precise sources of cohort selection effects observed here.

Secondly, this analysis is restricted to hippocampal volume within the ADNI dataset specifically; whether the same patterns of label inconsistency, technical drift, and survivor bias generalize to other brain regions and other longitudinal cohorts remains an open question for future work.

Despite these limitations, the consistency of the observed patterns across multiple analyses suggests that the identified sources of bias represent fundamental properties of the dataset rather than isolated artifacts. As such, the methodological considerations outlined in this work remain relevant for researchers developing machine learning models using ADNI and similar longitudinal neuroimaging resources.

### 4.6 Conclusion

The Alzheimer’s Disease Neuroimaging Initiative has become a foundational resource for machine learning applications in neuroimaging. However, its design as a longitudinal observational cohort introduces structural characteristics that complicate its use as a conventional labeled dataset.

In this study, we identified three key sources of bias within ADNI: variability between diagnostic labels and cognitive measurements, systematic differences in imaging features arising from evolving acquisition infrastructure, and cohort selection effects driven by longitudinal participant retention. These factors collectively influence both the inputs and targets used in machine learning models, challenging the assumption that ADNI provides stable and internally consistent data for predictive modeling.

Rather than representing isolated limitations, these findings highlight a broader methodological issue: the need to explicitly validate the assumptions underlying dataset usage in machine learning workflows. The graphical abstract summarizes each bias and its mitigation, providing a practical starting point for researchers using ADNI and similar longitudinal cohorts in machine learning.

By framing these considerations as integral components of the modeling process, this work provides a practical foundation for improving the robustness and interpretability of machine learning models trained on large-scale neuroimaging datasets. As machine learning becomes increasingly integrated into neuroimaging research, the validity of predictive models will depend not only on algorithmic sophistication, but also on careful examination of the datasets from which these models learn. ADNI remains an invaluable scientific resource, but its effective use requires explicit consideration of the longitudinal, technical, and diagnostic processes that shape the data itself.

## Acknowledgements

The preprinted version of this document can be found at…

## Author Contributions

PS – Conceptualization, Data Curation, Formal Analysis, Investigation, Methodology, Software, Writing & Reviewing (Original Draft)

HR – Writing (review & editing) & Original Draft

KR – Writing (review & editing) & Original Draft

AD – Writing (review & editing) & Original Draft

CD – Writing (review & editing) & Original Draft

MS – Project Administration, Supervision, Conceptualization, Writing (review & editing) & Original Draft

## Footnotes

1 ADNI4 is still ongoing and incomplete at the time of submitting this paper.

## References

Barnes, J., Bartlett, J.W., van de Pol, L.A. T Loy, C., Scahill, R.I., Frost, C., Thompson, P., Fox, N.C., 2009. A meta-analysis of hippocampal atrophy rates in Alzheimer’s disease. Neurobiology of Aging 30, 1711–1723. 10.1016/j.neurobiolaging.2008.01.010

Beer, J.C., Tustison, N.J., Cook, P.A., Davatzikos, C., Sheline, Y.I., Shinohara, R.T., Linn, K.A., 2020. Longitudinal ComBat: A method for harmonizing longitudinal multi-scanner imaging data. NeuroImage 220. 10.1016/j.neuroimage.2020.117129

Bigler, E.D., Skiles, M., Wade, B.S.C., Abildskov, T.J., Scheibel, R.S., Newsome, M.R., Mayer, A.R., Stone, J.R., Taylor, B.A., Tate, D.F., Walker, W.C., Levin, H.S., Wilde, E.A., 2020. FreeSurfer 5.3 versus 6.0: are volumes comparable? A Chronic Effects of Neurotrauma Consortium study. Brain Imaging and Behavior 14, 1318–1327. 10.1007/s11682-018-9994-x

Bron, E.E., Klein, S., Papma, J.M., Jiskoot, L.C., Venkatraghavan, V., Linders, J., Aalten, P., De Deyn, P.P., Jan Biessels, G., Classen, J.A., Smits, M., Niessen, W.J., van Swieten, J.C., van der Flier, W.M., Ramakers, I.H., van der Lugt, A., 2021. Cross-cohort generalizability of deep and conventional machine learning for MRI-based diagnosis and prediction of Alzheimer’s disease. NeuroImage: Clinical 31. 10.1016/j.nicl.2021.102712

Canevelli, M., Grande, G., Lacorte, E., Quarchioni, E., Cesari, M., Mariani, C., Bruno, G., Vanacore, N., 2016. Spontaneous Reversion of Mild Cognitive Impairment to Normal Cognition: A Systematic Review of Literature and Meta-Analysis. J Am Med Dir Assoc 17, 943–8. 10.1016/j.jamda.2016.06.020

Dalby, C., Dibble, A., Benini, S., Ferrari, D., Lyall, D.M., Harvey, M., Quinn, T., Muckli, L., Fracasso, A., Svanera, M., 2026. NeuroMorph: A Unified Morphological Reference Space for Cross-Disease Brain Profiling. 10.64898/2026.08.13.26359403

Dibble, A., Dalby, C., Sevegnani, M., Fracasso, A., Lyall, D.M., Harvey, M., Svanera, M., 2026. NeuroFM: Toward Precision Neuroimaging with Foundation Models for Individualized Brain Health Estimation. 10.64898/2026.03.27.26349489

Farina, M.P., Saenz, J., Crimmins, E.M., 2023. Does adding MRI and CSF-based biomarkers improve cognitive status classification based on cognitive performance questionnaires? 10.1371/journal.pone.0285220

Fischl, B., 2012. FreeSurfer. NeuroImage 62. 10.1016/j.neuroimage.2012.01.021

Fortin, J.-P., Cullen, N., Sheline, Y.I., Taylor, W.D., Aselcioglu, I., Cook, P.A., Adams, P., Cooper, C., Fava, M., McGrath, P.J., McInnis, M., Philips, M.L., Trivedi, M.H., Weissman, M.M., Shinohara, R.T., 2018. Harmonization of cortical thickness measurements across scanners and sites. NeuroImage 167, 104–120. 10.1016/j.neuroimage.2017.11.024

Fortin, J.-P., Parker, D., Tunc, B., Watanabe, T., Elliott, M.A., Ruparel, K., Roalf, D.R., Satterthwaite, T.D., Gur, R.E., Schultz, R.T., Verma, R., Shinohara, R.T., 2017. Harmonization of multi-site diffusion tensor imaging data. NeuroImage 161, 149–170. 10.1016/j.neuroimage.2017.08.047

Frenay, B., Verleysen, M., 2014. Classification in the Presence of Label Noise: A Survey. Transactions on Neural Networks and Learning Systems 25. 10.1109/TNNLS.2013.2292894

Holbrook, A.J., Tustison, N., Marquez, F., Roberts, J., Yassa, M.A., Gillen, D.L., 2020. Anterolateral entorhinal cortex thickness as a new biomarker for early detection of Alzheimer’s disease. Alzheimers Dement (Amst). 12. 10.1002/dad2.12068

Jack Jr, C.R., Knopman, D.S., Jagust, W.J., Petersen, R.C., Weiner, M.W., Aisen, P.S., Shaw, L.M., Vemuri, P., Wiste, H.J., Weigand, S.D., Lesnick, T.G., Pankratz, V.S., Donohue, M.C., Trojanowski, J.Q., 2013. Tracking pathophysiological processes in Alzheimer’s disease: an updated hypothetical model of dynamic biomarkers. Lancet Neurology 12, 207–16. 10.1016/S1474-4422(12)70291-0.

Johnson, W.E., Li, C., Rabinovic, A., 2007. Adjusting batch effects in microarray expression data using empirical Bayes methods. Biostatistics 8, 118–27. 10.1093/biostatistics/kxj037

Macdonald, K.E., Leung, K.K., Bartlett, J.W., Blair, M., Malone, I.B., Barnes, J., Ourselin, S., Fox, N.C., 2014. Automated Template-Based Hippocampal Segmentations from MRI: The Effects of 1.5T or 3T Field Strength on Accuracy. Neuroinformatics 12, 405–412. 10.1007/s12021-013-9217-y

Moreno-Torres, J.G., Raeder, T., Alaiz-Rodriguez, R., Chawla, N.V., Herrera, F., 2012. A unifying view on dataset shift in classification. Pattern Recognition 45, 521–530. 10.1016/j.patcog.2011.06.019

Petersen, R.C., Aisen, P.S., Beckett, L.A., Donohue, M.C., Gamst, A.C., Harvey, D.J., Jack jr, C.R., Jagust, W.J., Shaw, L.M., Toga, A.W., Trojanowski, J.Q., Weiner, M.W., 2010. Alzheimer’s Disease Neuroimaging Initative (ADNI). Neurology 74, 201–209. 10.1212/WNL.0b013e3181cb3e25

Pomponio, R., Erus, G., Habes, M., Doshi, J., Srinivasan, D., Mamourian, E., Bashyam, V., Nasrallah, I.M., Satterthwaite, T.D., Fan, Y., Launer, L.J., Masters, C.L., Maruff, P., Zhuo, C., Völzke, H., Johnson, S.C., Fripp, J., Koutsouleris, N., Wolf, D.H., Gur, Raquel, Gur, Ruben, Morris, J., Albert, M.S., Grabe, H.J., Resnick, S.M., Bryan, R.N., Wolk, D.A., Shinohara, R.T., Shou, H., Davatzikos, C., 2020. Harmonization of large MRI datasets for the analysis of brain imaging patterns throughout the lifespan. NeuroImage 208. 10.1016/j.neuroimage.2019.116450

Salthouse, T.A., 2014. Article Navigation Journal Article Selectivity of Attrition in Longitudinal Studies of Cognitive Functioning. The Journals of Gerontology Series B 69, 567–574. 10.1093/geronb/gbt046

Schmidt, M.F., Storrs, J.M., Freeman, K.B., Jack Jr, C.R., Turner, S.T., Griswold, M.E., Mosley Jr, T.H., 2018. A comparison of manual tracing and FreeSurfer for estimating hippocampal volume over the adult lifespan. Human Brain Mapping 39, 2283–2688. 10.1002/hbm.24017

Svanera, M., Savardi, M., Signoroni, A., Benini, S., Muckli, L., 2024. Fighting the scanner effect in brain MRI segmentation with a progressive level-of-detail network trained on multisite data. Medical Image Analysis 93. 10.1016/j.media.2024.103090

Zheng, B., Udeh-Momoh, C., Watermeyer, T., de Jager Loots, C.A., Ford, J.K., Robb, C.E., Giannakopoulou, P., Ahmadi-Abhari, S., Baker, S., Novak, G.P., Price, G., Middleton, L.T., 2022. Practice Effect of Repeated Cognitive Tests Among Older Adults: Associations With Brain Amyloid Pathology and Other Influencing Factors. Frontiers Aging Neuroscience 14. 10.3389/fnagi.2022.909614

